# Revealing druggable cryptic pockets in the Nsp-1 of SARS-CoV-2 and other β-coronaviruses by simulations and crystallography

**DOI:** 10.1101/2022.05.20.492819

**Authors:** Alberto Borsatto, Obaeda Akkad, Ioannis Galdadas, Shumeng Ma, Shymaa Damfo, Shozeb Haider, Frank Kozielski, Carolina Estarellas, Francesco Luigi Gervasio

## Abstract

Non-structural protein 1 (Nsp1) is a main pathogenicity factor of *α*- and *β*-coronaviruses. Nsp1 of severe acute respiratory syndrome coronavirus 2 (SARS-CoV-2) suppresses the host gene expression by sterically blocking 40S host ribosomal subunits and promoting host mRNA degradation. This mechanism leads to the downregulation of the translation-mediated innate immune response in host cells, ultimately mediating the observed immune evasion capabilities of SARS-CoV-2. Here, by combining extensive Molecular Dynamics simulations, fragment screening and crystallography, we reveal druggable pockets in Nsp1. Structural and computational solvent mapping analyses indicate the partial crypticity of these newly discovered and druggable binding sites. The results of fragment-based screening via X-ray crystallography confirm the druggability of the major pocket of Nsp1. Finally, we show how the targeting of this pocket could disrupt the Nsp1-mRNA complex and open a novel avenue to design new inhibitors for other Nsp1s present in homologous *β*-coronaviruses.

## Introduction

Coronaviruses (CoVs) are the largest family of RNA viruses identified to date. CoVs are members of the subfamily *Coronavirinae* classified into four genera *α*-, *β*-, *γ*-, and *δ*-*coronavirus*. Common human CoVs belong to the first two genera and include the 229E (*α*-), NL63 (*α*-), OC43 (*β*-), and HKU1 (*β*-). SARS-CoV-2, which led to the COVID-19 pandemic declared in March of 2020 as the severe acute respiratory syndrome-related coronavirus, SARS-CoV-1, and the Middle East respiratory syndrome-related coronavirus, MERS-CoV, all belong to the *β*-coronavirus genera and are suggested to originate from bats.^1–3^ The genome of different CoVs typically encodes four structural proteins, namely spike, envelope, membrane and nucleocapsid, and two large polyproteins, pp1a and pp1ab, that are later cleaved into several non-structural proteins.^4^ Due to the SARS-CoV-2 pandemic, significant efforts have been directed to the study and inhibition of these proteins such as the spike protein and the protease M^pro^.^5^ Among the different proteins that are involved in the pathogenicity of SARS-CoV-2 is the Non-Structural Protein 1 (Nsp1), a small 180 residue protein^6^ whose function has been studied comparatively less than the rest of the pathogenic proteins. The structure of Nsp1 can be divided into three domains, a structured core, and two disordered domains corresponding to the N- and C-termini of the protein. The structured domain is composed of 117 residues (Glu10-Asn126) and displays an *α/β*-fold. The disordered C-terminal domain (Gly127-Gly180) was shown to partially fold into a helix-loop-helix pattern when bound to the 40S ribosomal subunit (Figure 1).^7^

**Figure 1.**
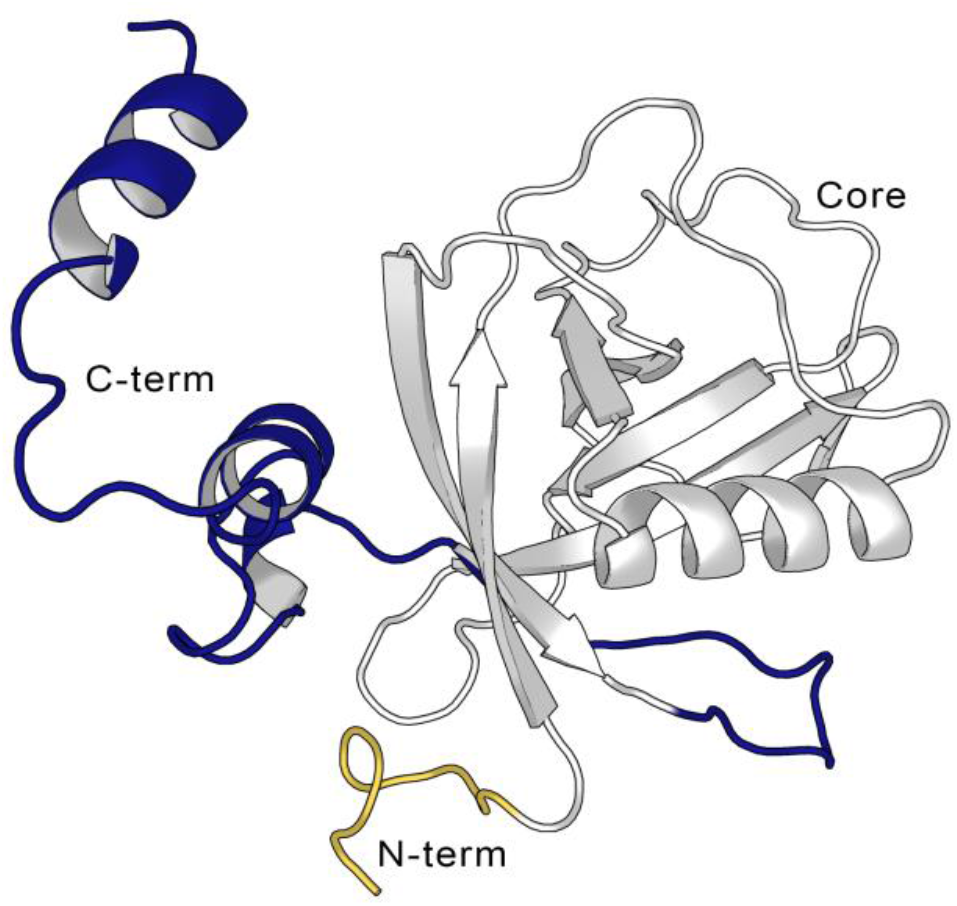
Cartoon representation of the full-length Nsp1 structure from the AlphaFold^8,9^ model, showing the N-terminus (in yellow, aa Met1-Asn9), the Nsp1_N_ core (in grey, aa Glu10-Asn126), and the C-terminus (blue, aa Gly127-Gly180).

It is worth noting that Nsp1 is only present in the *α*- and *β*-, but not in the *γ*-, and *δ-CoVs*. The structural analysis of Nsp1_N_ structures from *α*-(PDB entry 3ZBD and 5XBC) and *β*-(PDB entry 2HSX) CoVs suggests that, despite the low sequence homology, the Nsp1 of the two genera displays a high structural similarity, with RMSD values ranging from 1.8 to 2.4 A.^10,11^ The structural similarity of the *α*-CoVs and *β*-CoVs Nsp1 is reflected by their biological function as both are involved in the regulation of the host and viral gene expression. Multiple studies have shown that the expression of Nsp1 inhibits the translation in host cells by a combination of different mechanisms. For instance, Nsp1 sterically blocks the mRNA tunnel in the 40S ribosomal subunit,^7^ the 43S preinitiation complex and the non-translating 80S ribosomes.^7,12,13^ Moreover, it has been shown that Nsp1 can also trigger the cleavage of the host mRNA and hinder its nuclear export to the cytosol.^14^ These mechanisms concur in downregulating the translation-mediated innate immune response of the host cell, hence mediating the observed immune evasion capabilities of SARS-CoV-2.^12,15^ Interestingly, only host mRNAs are subjected to Nsp1-mediated endonucleolytic cleavage whereas viral mRNAs escape from this translation suppression mechanism. Experimental results have shown that the interaction of N-terminal Nsp1 with the stem loop 1 (SL1) at the viral mRNA 5’ UTR region is necessary to avoid the Nsp1-mediated translation shutdown and cleavage in infected cells.^16,17^

The regulatory role of Nsp1 in the viral replication and gene expression has also been demonstrated by mutations in the Nsp1 coding region of the transmissible gastroenteritis virus (TGEV, an *α*-CoV infecting pigs) and the murine hepatitis virus (MHV, a *β*-CoV infecting mice) genomes. According to these mutagenesis studies, blocking the function of Nsp1 in different viruses leads to a drastic reduction or elimination of the infectious virus.^18^

Altogether, the high structural similarity across different *α*- and *β*-CoVs from different organisms, the fact that Nsp1 has no homologues outside of the CoVs, as well as its crucial role in mediating immune evasion, make Nsp1 a valuable target for developing antiviral drugs, not only for the ongoing COVID-19 pandemic but also, to prevent future pandemic outbreaks caused by new variants. However, the largest folded domain of Nsp1, namely the N-terminal core region (Nsp1_N_), corresponds to a small compact domain that shows predominantly small superficial cavities, which complicates rational drug design efforts. In spite of Nsp1 being a validated target for therapeutic intervention, very few studies explored Nsp1 for structure-based drug discovery, and only one of them reported the binding of a ligand to the C-terminal domain of SARS-CoV-2 Nsp1.^19^ To date, no ligand-bound Nsp1 crystal structures are available, making the ones presented here and in another study from our group the first fragment-bound SARS-CoV-2 Nsp1 crystal structures to date.

Here, we used a combination of computational and experimental approaches to explore the druggability of Nsp1 in SARS-CoV-2, and the possibility to expand the findings to homologous Nsp1s. In particular, we have used modelling, enhanced sampling simulations, virtual screening, fragment soaking, and X-ray crystallography to identify druggable binding pockets, including hidden (cryptic) ones, and evaluate the potential of ligands to interfere with the formation of Nsp1-RNA complexes. Our enhanced sampling simulations predict four partially cryptic binding pockets of which one is validated by crystallography. Moreover, taking into consideration the 3D similarity between different Nsp1s in *α*- and *β*-CoVs, we have extended our analysis to assess the conservation of the pockets across various Nsp1s of *α*- and *β-CoVs* infecting humans. The results of this research can be used as a stepping stone for the design of Nsp1 inhibitors for SARS-CoV-2 and, potentially for other *α*- and *β*-CoVs.

## Results and discussion

### Structural assessment of Nsp1 pockets

To the date of writing this paper, the only available X-ray structure of apo SARS-CoV-2 Nsp1 contains only the N-terminal region (aa 10 −126 PDB entry 7K7P)^20^, hereafter referred to as Nsp1_N_ system. Starting from this crystal structure, we have characterized the possible binding pockets present in the 3D structure by means of pocket detection algorithms, namely DoGSiteScorer^21^ (available via the ProteinPlus webserver) and Fpocket^22^, showing similar results (Figure 2 and Figure 2 - Figure supplement 1). The analysis indicates the presence of some putative binding sites on the protein surface. Specifically, two different areas were identified to harbor potential binding pockets by both algorithms. The first one is sandwiched between the entrance of the *β*-barrel and the *α*-helix displaying a more evident pocket-like structure with a concave topology (Figure 2, left). The second region, located on the opposite side of the β-barrel, is characterized by a groove-like topology and spans a larger area on the protein surface but is very shallow (Figure 2, right).

**Figure 2.**
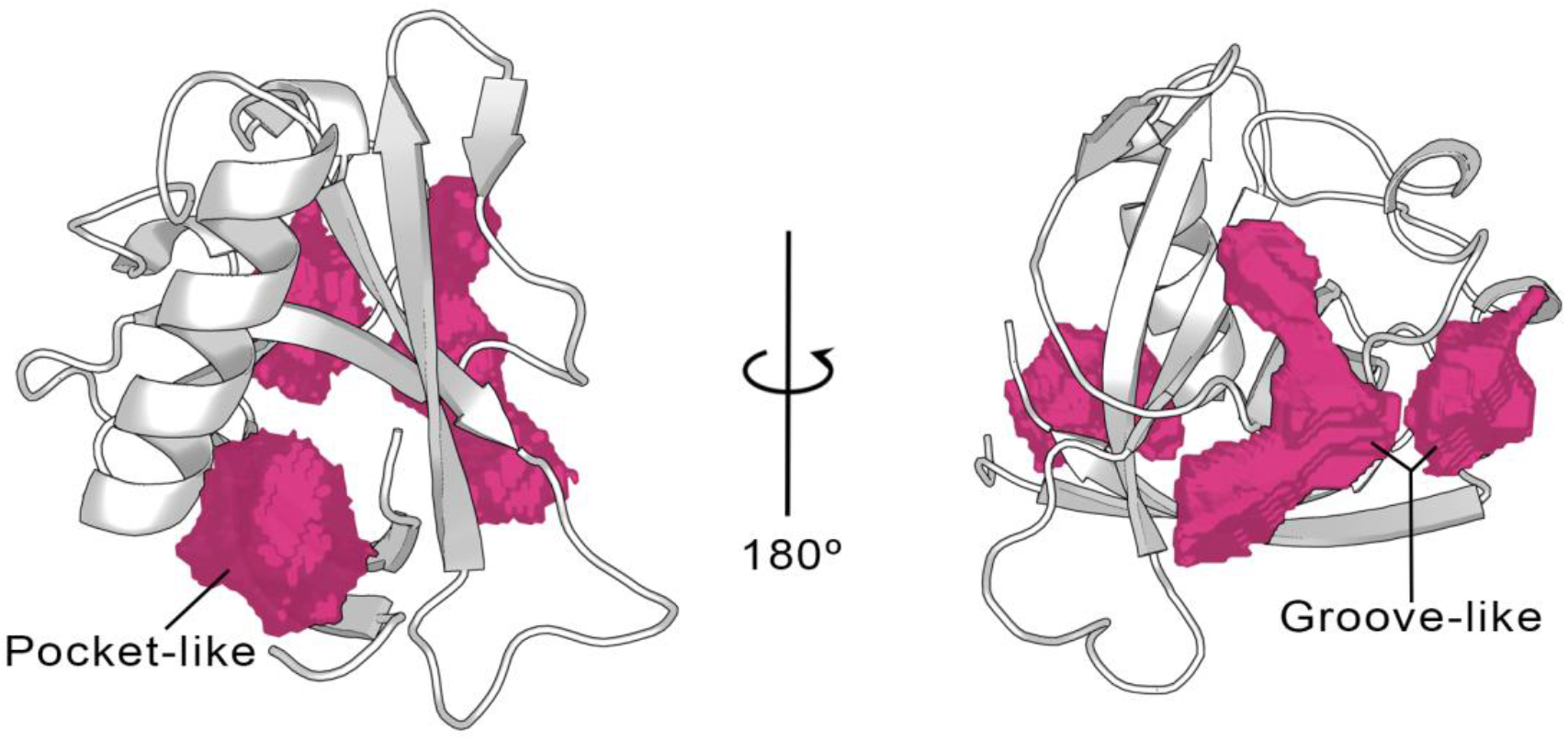
Cavities identified on the Nsp1_N_ crystal structure (PDB entry 7K7P) by the ProteinPlus server for the concave pocket-like structure between the β-barrel and the α-helix and the groove-like topology.

To investigate more in-depth the nature of the pocket-like structure and the possible opening of hidden (cryptic) binding sites in the grooved region, we have carried out a 1 μs long unbiased Molecular Dynamics (MD) simulation of Nsp1_N_ in its apo state. The timeseries of the root-mean-square deviation (RMSD) shows that the system remains stable throughout the simulation (Figure 3–figure supplement 1). The analysis of the pockets observed along the MD simulation through the MDpocket22 program confirms the presence of a pocket-like structure, hereafter referred to as pocket 1, at the same place as the one predicted for the X-ray structure by ProteinPlus and Fpocket. A detailed analysis of pocket 1 indicates that the residues forming this pocket are mostly located between the N- and C-termini of the Nsp1_N_ protein, namely between the α-helix and the two *β*-strands of the *β*-barrel (Figure 3A, left). The analysis of the pocket’s volume confirms the stability of pocket 1, with an average value of 410 ± 150 Å^3^ (Figure 3B). Interestingly, the volume obtained during the simulation is significantly larger than the one predicted based on the crystal structure (217 Å^3^), since during the simulation the structural features of any pocket are subject to fluctuations due to the rearrangements of the amino acid side chains. However, in this case, the difference in volume between the crystal structure and simulations is the result not only of the residue fluctuations, but also of the different residues identified around the pocket. Specifically, the residues close to the *β*-barrel form a larger cavity in the simulation than the one observed in the Nsp1_N_ crystal structure. This finding suggests that pocket 1 is partially cryptic. It is worth noting that some of these residues are located at the beginning of the C-terminus loop of the protein, suggesting that the binding of a ligand to this region can potentially affect the dynamics of the Nsp1 C-terminus, and possibly release the blockage of host RNA entry into ribosome for translation. Additionally, as pocket 1 is close to the rival RNA binding region,^23^ the binding of a ligand in this pocket could also interfere with the interaction with SL1 of viral RNA, leading to the failure in evasion of translation shutdown and cleavage of viral RNA.The second pocket identified during the MD simulations, namely pocket 2, is located in the grooved region. The structural analysis of pocket 2 over the simulation does not present significant differences with respect to the X-ray structure. In this pocket, the residues are mostly distributed between the loops connecting different *β*-strands of the *β*-barrel and the *α*-helix (Figure 3A, right). Likewise, the volume calculated during the simulation is 240 ± 114 Å^3^, in agreement with the volume predicted by ProteinPlus for the Nsp1_N_ crystal structure (150 Å^3^).

**Figure 3.**
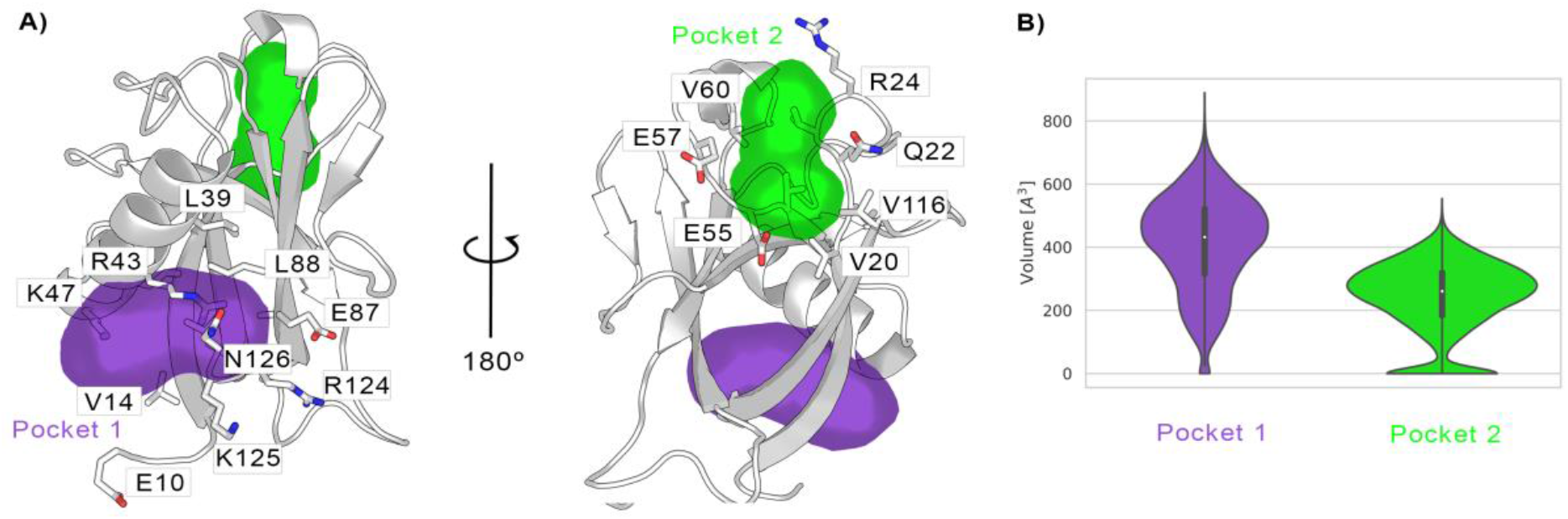
A) Cavities identified on Nsp1_N_ along the 1 μs unbiased MD simulation, namely pocket 1 (purple), and pocket 2 (green), with the main residues displayed in sticks. B) The volume distribution of each pocket along the unbiased MD simulations.

### Crypticity assessment of Nsp1 pockets

Encouraged by the results obtained from the unbiased MD simulations, we used SWISH (Sampling Water Interfaces through Scaled Hamiltonians) with mixed solvents, an enhanced-sampling method developed by our group to explore cryptic binding pockets. SWISH is a Hamiltonian replica-exchange method that improves the sampling of hydrophobic cavities by scaling the interactions between water molecules and protein atoms. It has been shown to be very effective in sampling the opening of hidden (cryptic) cavities in several different targets.^24^ We have run six replicas of 500 ns each, considering a concentration of 1 M benzene as co-solvent, the presence of which is expected to stabilize any pocket that will open transiently during the simulations (Figure 4 - figure supplement 1). The resulting trajectories have been analyzed with MDpocket.^22^ It is worth noting that, in addition to confirming the presence of pocket 1 and pocket 2, our analysis shows the presence of two new pockets, namely, pocket 3 and pocket 4. These two pockets are both located on the exterior of the *β*-barrel, proximal to each other and near pocket 2 (Figure 4 A). Most of the residues composing pocket 3 are part of the *β*-sheets of the *β*-barrel whereas most of the residues in pocket 4 are part of the loops connecting different *β*-sheets of the barrel. Interestingly, our previous analysis performed over the X-ray structure identified a shallow groove on the surface of Nsp1 connecting pockets 2, 3 and 4. However, neither of the two pocket detection algorithms used on the X-ray structure detected any evident deep cavity in this region. On the contrary, the analysis of the resulting SWISH trajectories clearly shows the presence of two distinct cavities with deep pocket-like structures. The absence of these sites in the crystal structure of Nsp1_N_ and the fact that these cavities remained closed during the MD simulations indicate the cryptic nature of pocket 3 and pocket 4.

**Figure 4.**
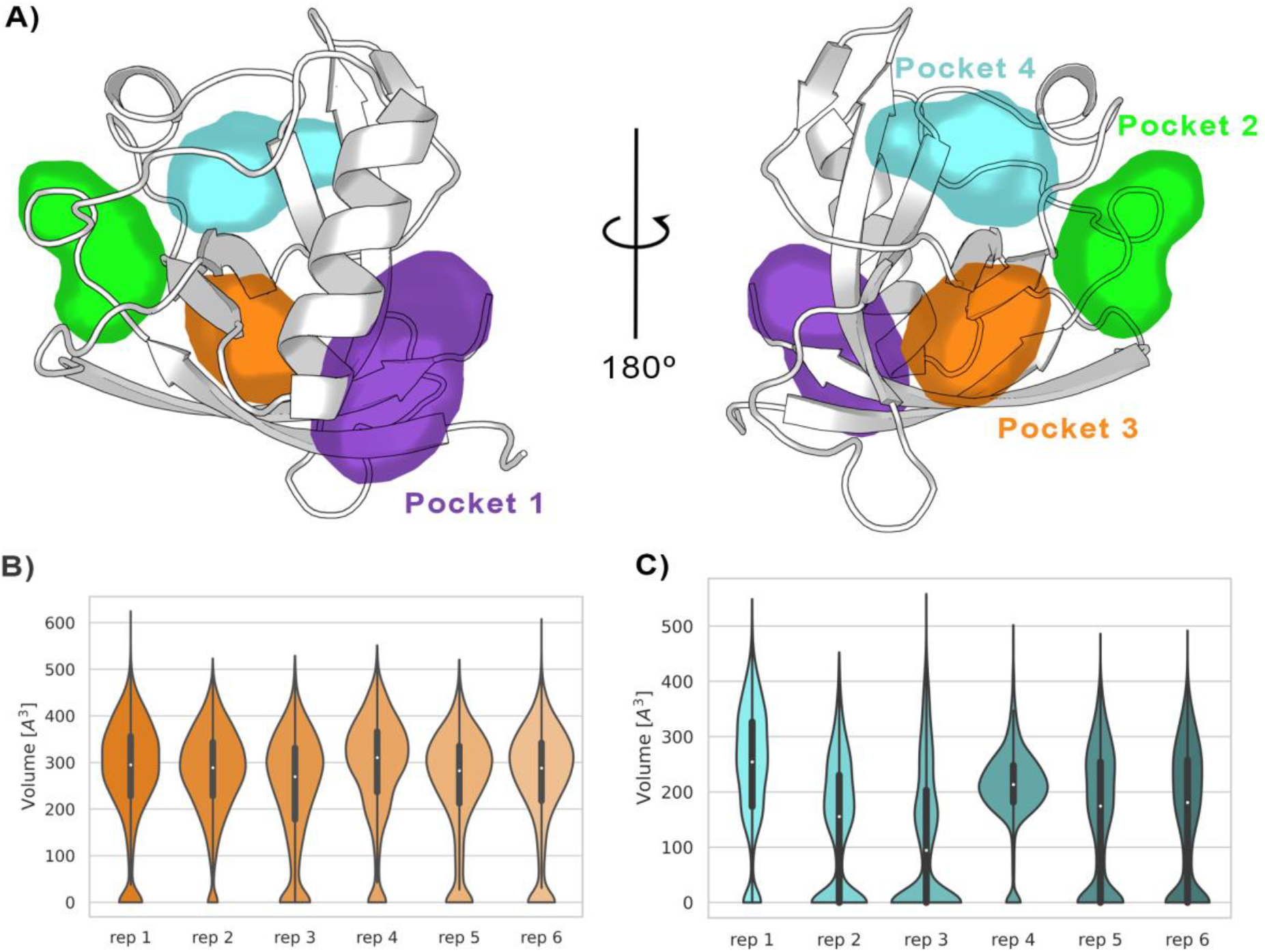
(A) Pockets sampled during the 500 ns of replica-exchange SWISH simulations. Volume distributions of the cryptic binding sites pocket 3 (B) and pocket 4 (C) along the six replicas of the SWISH simulations.

The average volume of pocket 1 during the SWISH simulations (471 ± 128 Å^3^) is comparable to the one obtained in the MD simulations (410 ± 150 Å^3^) for the same pocket (Figure 4 - Figure supplement 2A). The topology of pocket 1, i.e., the residues identified around the pocket, is the same for the MD and SWISH simulations (Figure 3). Likewise, the volume analysis of pocket 2 also indicates that this pocket remains open, and its volume is essentially equal to the one sampled during the unbiased simulation (214 ± 113 Å^3^, Figure 4 - Figure supplement 2B). More interestingly, the opening of pocket 3 (268 ± 115 Å^3^) during the SWISH simulations revealed a pocket of comparable size to pocket 2. The time series of the volume profiles indicates that the pocket starts from a closed-like conformation that opens quickly to a more pocket-like conformation. Once opened, it maintains this open-like conformation in all the replicas (Figure 4B). A different behavior was observed for pocket 4. This pocket, the smallest one sampled during the SWISH simulations (169 ± 120 Å^3^), displays diverse volume profiles across most SWISH replicas. Nonetheless, we were able to sample an open-like conformation for this pocket in at least two of the six replicas, suggesting that a better combination of molecular probes or higher λ factors could improve the sampling of this cryptic site. Interestingly, as suggested by the preliminary analysis of the X-ray structure, the simulations suggest as well that the shallow groove most likely has a key structural role connecting three cryptic pockets of the *β*-barrel region of the Nsp1_N_, spanning this area of the protein. Our findings highlight the potential of this region to be exploited in Fragment-Based Drug Design to design larger ligands that could bind to different combinations of these pockets, increasing the specificity of the individual pockets.

To further investigate the structural features of the predicted pockets, we analyzed the unbiased MD simulations with FTDyn and FTMap programs.^25^ These programs have proven to be accurate in locating binding hot spots in proteins. It is based on the fast and accurate distributions of 16 different small organic probes docked and mapped onto the protein surface, retaining the lowest energy probes, which are finally clustered. The clusters obtained for the different probes are clustered together to generate a final consensus cluster, from which the main binding hot spots are then identified. We started by employing the FTDyn webserver, a faster version of the FTMap algorithm without local minimization, to identify the most probable conformation able to bind fragments. We extracted 25 representative conformations from the MD simulation of Nsp1_N_ and determined the median number of interactions between the probes and each protein residue across the conformations selected. This step allows us to identify the most likely binding site residues, that is, the residues with the highest number of probe contacts. From the initially selected conformations, we have retained only 11 structures that present higher-than-average probe-residue contacts. Subsequently, the 11 structures retained were post-processed with FTMap to improve the prediction of binding host spots on Nsp1_N_ surface (see details in Methods Section and Figure 5-Table supplement 1).

Our results are in good agreement with the previous analysis, regarding the location of the binding sites found for 107 consensus clusters obtained for Nsp1_N_. In pocket 1, 50 out of the 107 consensus clusters are mapped into it, most of them corresponding to protein-fragment complexes with the highest binding energy (Figure 5A and Figure 5-Table supplement 1). Interestingly, the clusters are distributed across the entire MD- and SWISH-sampled volume of pocket 1 and are not only limited to the small region observed in the X-ray structure, suggesting that pocket 1 is partially cryptic. The computational mapping also confirmed the presence of 50 consensus clusters in the vicinity of pockets 2, 3 and 4 (Figure 5B). In all selected bound-like Nsp1_N_ structures, we identified at least two probes consensus clusters located in the region corresponding to the two cryptic sites previously identified by our SWISH simulations. In some cases, the second-best binding hotspot is in the presumably cryptic region, suggesting the ability of this region to accommodate molecular fragments (Figure 5-Table supplement 1).^26^ Overall, this analysis strongly suggests that the main binding sites on Nsp1_N_ are in the region corresponding to the identified cryptic pockets and highlights their potential use for fragment binding.

**Figure 5.**
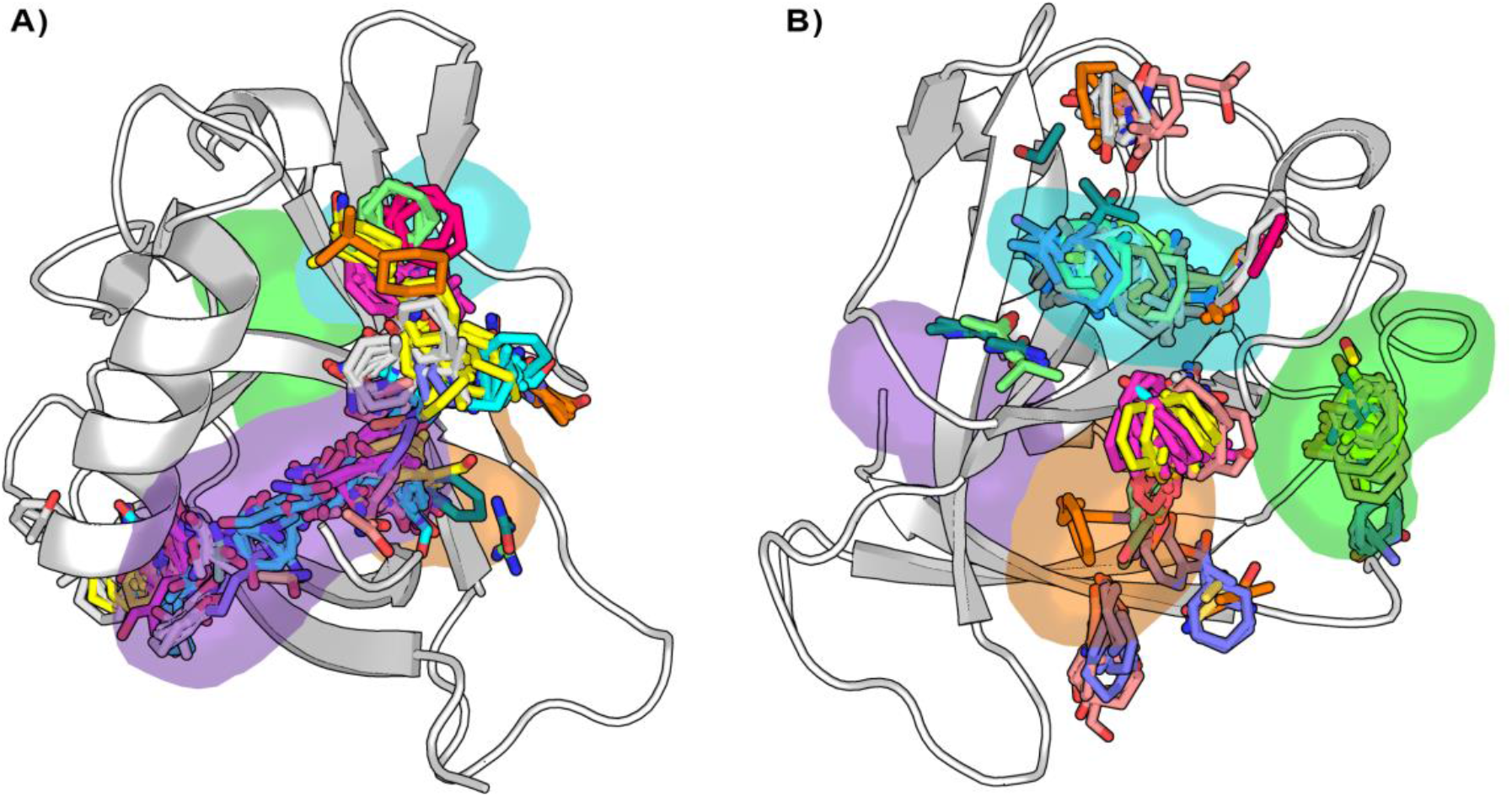
Distribution of the binding hot spots on the Nsp1_N_ surface around A) pocket 1 and B) pockets 2, 3 and 4. Multiple consensus clusters are shown in sticks. Each consensus cluster is represented in different colors.

### Crystallographic confirmation of Nsp1 cryptic pockets

To validate our computational findings, we proceeded with the soaking of Nsp1_N_ crystals with 60 potential fragment hits obtained by computational methods. Nsp1_N_ has been crystallized containing only the structured domain, i.e., the domain from Glu10 to Asn126, which displays the characteristic α/β fold (see Methods for details). This structure was used for crystal soaking experiments in which 60 distinct fragment hits, obtained from the Maybridge Ro3 library, were tested and validated through X-ray diffraction experiments using the Pan-Dataset Density Analysis (PanDDA) method, developed to analyze the data obtained from crystallographic fragment screenings.^27^ Of these 60 fragments, one fragment was found in pocket 1 as previously identified in our simulations. Data collection and refinement statistics are summarized in Fig 6-Table supplement 1. The asymmetric unit contains one molecule of Nsp1. The model covers the sequence from Glu10 to Asn126 (E10 to N126). The chemical structure of the fragment hit is shown in Figure 6B. To study the fragment hit using orthogonal biophysical assays, we employed microscale thermophoresis (MST) and thermal shift (TSA) assays. 2E10 binds to Nsp1 pocket 1 with a Kd value of 2.74 ± 2.63 mM, indicating rather good binding for a fragment. In contrast we did not observe any stabilization of Nsp1 by the fragment. The fragment obeys the rule of 3 (Ro3) with a molecular mass of 172.9 Da, a calculated MolLogP of 2.66 and a total of 3 hydrogen bond donors and acceptors. Fragment hit 2E10 combines two fused 5- and 6-membered ring systems containing one acetamide substituent at the phenyl group A range of residues located in binding pocket 1 including Glu10, Val14, Arg43, Leu46, Lys47 and Leu123, establish hydrophobic interactions with the ligand. (Figure 6C). The acetamide substituent establishes a hydrogen bond interaction with the side chain of Lys125.

**Figure 6.**
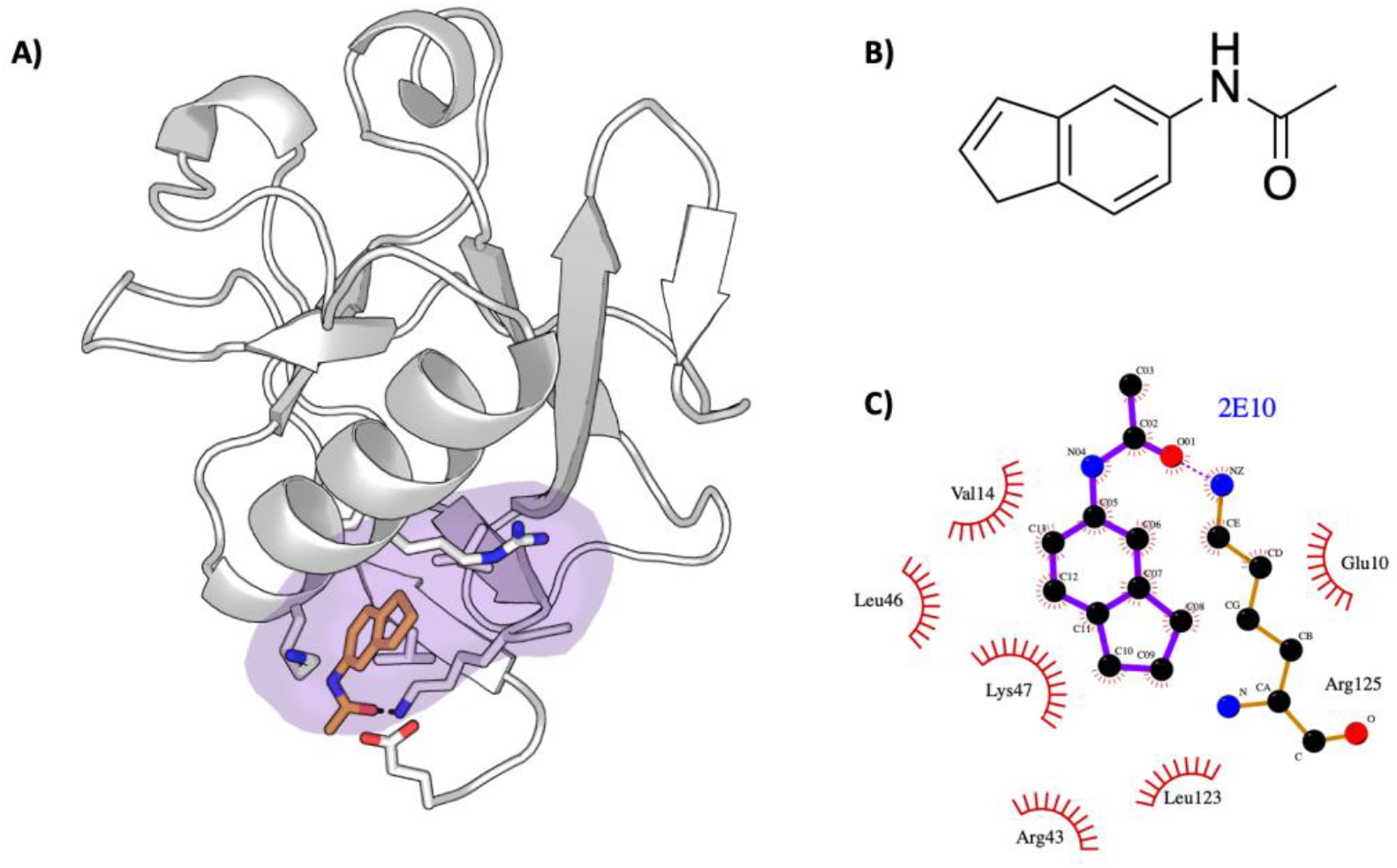
Characterization of the SARS-CoV-2 Nsp1-2E10 complex. A) Binding pose of the fragment hit obtained by crystal soaking and structure determination methods. The fragment is located in pocket 1. B) Chemical structure and name of fragment hit. C) Magnification of the Nsp1_N_ −2E10 binding pocket showing the interactions the fragment establishes with residues of Nsp1. Hydrophobic interactions are shown by red half-moons and the hydrogen bond interaction is displayed with a dotted purple line.

Although, at first, this may seem like a very low number of hits, a detailed analysis of the crystal packing provides an explanation. Figure 7A shows the central Nsp1_N_ structure (white) with the pockets obtained from our SWISH simulations, together with the neighboring Nsp1 structures that interact directly with pocket 1 in the full crystal packing (dark grey). Evidently, the central part of the pocket is completely accessible for fragment binding during the crystal soaking experiments. However, the grey structures are occupying the main hotspots identified by our previous FTMap analysis (Figure 5A) impeding its ability to bind more suitable fragments in pocket 1. Likewise, Figure 7B shows the central Nsp1 structure in cartoon representation (white) with the pockets obtained from our SWISH simulations, along with the neighboring Nsp1 structures directly interacting with pockets 2, 3, and 4 in the full crystal packing (dark grey). The red regions highlighted in Figure 7 involve direct contacts made with the pocket environment. It is worth emphasizing that, in this case, these direct contacts involve regions previously determined by the FTMap analysis as potential and key hotspots for the three cryptic pockets identified by SWISH. Taken altogether, soaking Nsp1_N_ crystals with fragments presents some limitations to implementing an effective computational fragment screening of the Nsp1. Nevertheless, we would like to stress that these results provide a possible explanation for why we did not obtain more fragments crystallized in the positions of the identified cryptic pockets. Moreover, the crystallization of a single fragment in pocket 1 is an encouraging sign of the reliability and efficacy of our computational techniques for cryptic binding pocket detection.

**Figure 7.**
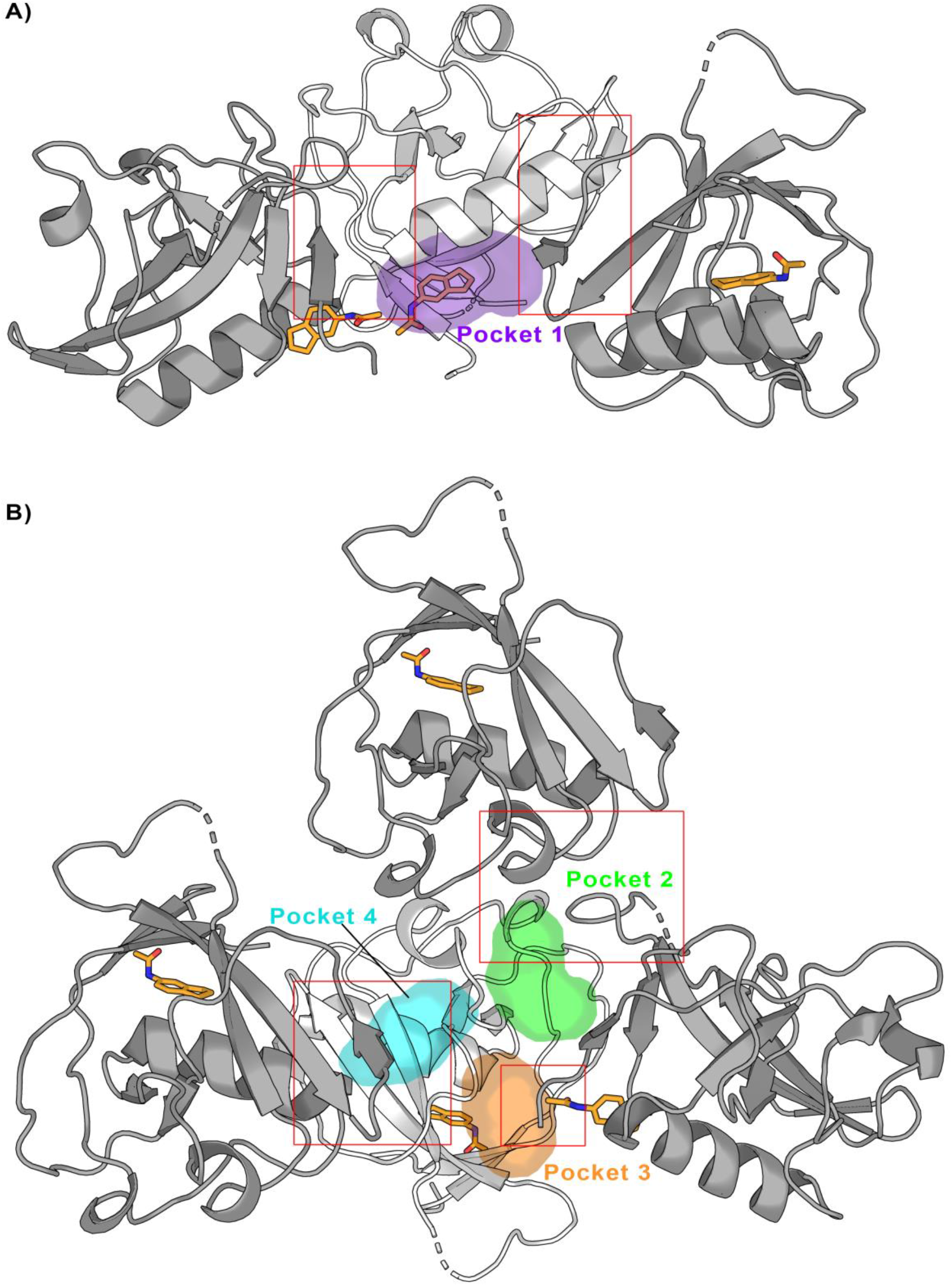
Crystal packing of the Nsp1_N_-fragment complex (code 2E10) was obtained from X-ray soaking experiments. The direct crystal contacts around A) pocket 1 and B) pockets 2, 3 and 4 are highlighted with squares.

### Disrupting Nsp1-RNA complex by means of cryptic pockets

The interaction of Nsp1 with viral RNA can release the inhibition of the viral RNA translation in host cells via the combination of different mechanisms, favoring the survival of the virus. Therefore, the disruption of the Nsp1-RNA complex can affect the life cycle of the virus. With this idea in mind, and taking into consideration the pockets found in Nsp1_N_, we asked whether the binding of fragments to pocket 1 could hinder the Nsp1-RNA complex formation. To this end, we have modelled two Nsp1-RNA complexes, testing their stability through MD simulations. It has been shown experimentally that the first stem loop (SL1), comprising nucleotides 7-33 of the 5’ UTR region of SARS-CoV-1 and SARS-CoV-2 RNA, is the one that interacts with Nsp1.^15–17,28^ Therefore, we started by modelling the 3D structure of SL1 RNA. Regarding the Nsp1, we have considered the model of the full-length Nsp1 obtained by AlphaFold^8,9^, named Nsp1_FL_. Using the Nsp1_FL_ and SL1 models, we have run protein-RNA docking using the HADDOCK program.^29,30^ The analysis of the docked Nsp1_FL_-RNA complexes suggests that two different protein-RNA complexes, named model A and model B, well capture the experimentally validated contacts between the Nsp1_FL_ and SL1. The Nsp1_FL_ in models A and B binds to diametrically opposite positions on the SL1, with Nsp1 in model A interacting predominately with the groove of SL1 (Figure 8A) and in model B with its backbone (Figure 8 - Figure supplement 1). Since there was no structural or experimental reason to exclude one over the other, we assessed their stability over the course of a 500 ns long MD simulation. The root-mean square deviation (RMSD) of the two Nsp1_FL_-RNA complexes fluctuate between 5.5 and 6.5 Å for models A and B respectively (Figure 8 - Figure supplement 2). However, a more detailed RMSD analysis of each component of the two protein-RNA systems reveals that Nsp1_FL_ is more stable in model A than in model B, with an average RMSD of 5.0 and 7.0 Å respectively. This result suggests that the Nsp1_FL_ in model A displays a more stable interaction with RNA. However, the observed RMSD difference alone between the two models is not sufficient to let us propose model A over B as the most probable mode of Nsp1-RNA interaction (Figure 8 - Figure supplement 2).

**Figure 8.**
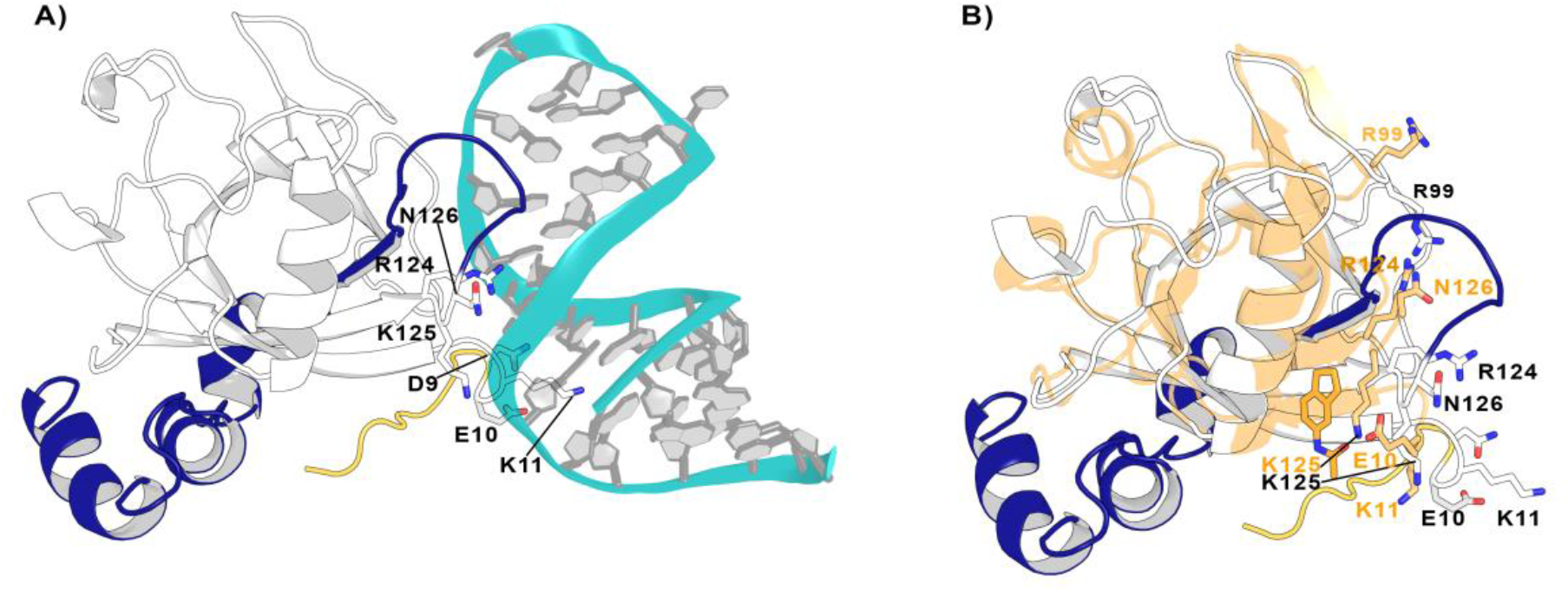
Nsp1_FL_-RNA complex obtained from the HADDOCK program. A) Residues in the proximity of pocket 1 involved in crucial Nsp1_FL_-SL1 contacts are displayed in sticks. B) Crystal structure from soaking experiments (orange transparent cartoon) is superimposed over the Nsp1_FL_ of the Nsp1_FL_-RNA model. The most important residues for the Nsp1_FL_-RNA interaction are highlighted in white (Nsp1_FL_ model) and orange (PDB entry 2E10 from our soaking experiments) sticks. Figures obtained from model A.

Since both models not only capture the important interactions between Nsp1_FL_ and viral RNA described experimentally but are also stable over the course of the MD simulations, we used them to verify if the previously identified pockets could be exploited to impede the Nsp1_FL_-RNA complex. Interestingly, out of the four pockets in Nsp1_N_, pocket 1 is positioned close to the Nsp1_FL_-RNA interface. More specifically, some residues of the C-terminal loop of Nsp1_FL_ around pocket 1 interact with the viral RNA, namely Arg124, Lys125 and Asn126. Moreover, Glu10 is oriented towards pocket 1 and both Asp9 and Lys11 establish contacts with the SL1 moiety (Figure 8A). Therefore, these results suggest that pocket 1 can be a crucial target for rational drug design since the residues that define this pocket are directly interacting with the viral RNA and the binding of a ligand to this pocket can impair the Nsp1_FL_-RNA interaction.

To further investigate the effect of ligand binding to pocket 1 and how this would affect the Nsp1-SL1 interface, we superimposed the crystal structure from the soaking experiments on the Nsp1_FL_-RNA model A (Figure 8B). Interestingly, in the Nsp1_FL_-RNA complex, some residues of the N-terminus (Asp9, Glu10 and Lys11) and the core (Arg99, Arg124, Lys125 and Asn126) of Nsp1_FL_ are directly interacting with the SL1 of RNA. When compared with the structure obtained from soaking experiments (Glu10 to Asn126), some of these residues, specifically Glu10 and Lys11 are pointing towards the fragment, Arg99 are pointing to the surface, and Arg124 and Asn126 are displaced due to the ligand binding, all of them changing the orientation observed in the Nsp1_FL_-RNA complex. In the orientation observed in the structure obtained from soaking experiments, these residues could not interact with the RNA. As these residues are essential for the Nsp1-RNA interaction in SARS-CoVs, therefore the binding of a ligand in pocket 1 could diminish the Nsp1-RNA interaction.^16,31^ Additionally, a recent study where site-directed mutagenesis was performed on the Nsp1 N-terminal and core region (Nsp1_N_) has demonstrated the functional role of Arg99, Arg124 and Lys125 residues in host expression shutdown, since the mutation of these residues to Ala compromised the binding of Nsp1 to the host 40S ribosomal subunit and increased the dissociation constants with purified ribosomes.^32^ Therefore, these findings support the hypothesis that a disruption of the interactions between these residues and SL1 by the presence of a fragment would lead to a decrease in virus-induced host shutoff.

### The conservation of Nsp1 in different CoV genera

Given the identified cryptic pockets on the Nsp1 of SARS-CoV-2 and their implication in the interaction with the RNA, we sought to analyze thoroughly the structural similarity of the Nsp1 among different CoVs. We asked whether a drug designed for Nsp1 of SARS-CoV-2 could also be useful to inhibit Nsp1 of other CoVs. As mentioned in the introduction, of the four CoV genera, only *α*- and *β*- are common human CoVs, and Nsp1 is expressed only in these two genera. To demonstrate the homology relationship of the Nsp1_N_ domain within different viruses belonging to the *α*- and *β*-genera, we have performed a phylogenetic analysis considering this domain, for both *α*- and *β*-CoV genera from the Conserved Domain Database (CDD) hosted at the National Center for Biotechnology Information (NCBI).^33^ The analysis of the 283 Nsp1_N_ sequences available until the end of February 2022 shows the distribution of Nsp1 homologues for different *α*-(TGEV- and PDEV-like) and *β*-(MERS-, HKU9-, SARS-, and MHV-like) CoVs (Figure 9A). In particular, we have performed a pair sequence alignment considering one representative Nsp1 protein from each of the *α*- and *β*-CoVs sub-families presented in our phylogenetic tree. ^34^ The alignment demonstrates that the sequence identity varies widely depending on the homologue proteins under consideration (Figure 9-Table supplement 1). Subsequently, to decrease the high heterogeneity, we have considered the Nsp1_N_ sequence from SARS-CoV-2 and three high identity homologues corresponding to Nsp1 proteins from the human SARS-CoV-1, and two Bat CoVs, namely BatCoV HKU3 and RaTG13 (NCB1 accession numbers: MT782115.1 and MN996532.2, respectively).^11^ The selected homologues share a high sequence identity with SARS-CoV-2 Nsp1 ranging from 86% to 93%. Figure 9B shows this high sequence identity between CoVs from different organisms, especially of the residues surrounding the cryptic pockets found in Nsp1 from SARS-CoV-2. Our analysis shows that for the four sequences, all the important residues of the pockets are conserved in the selected *β*-CoVs.

**Figure 9.**
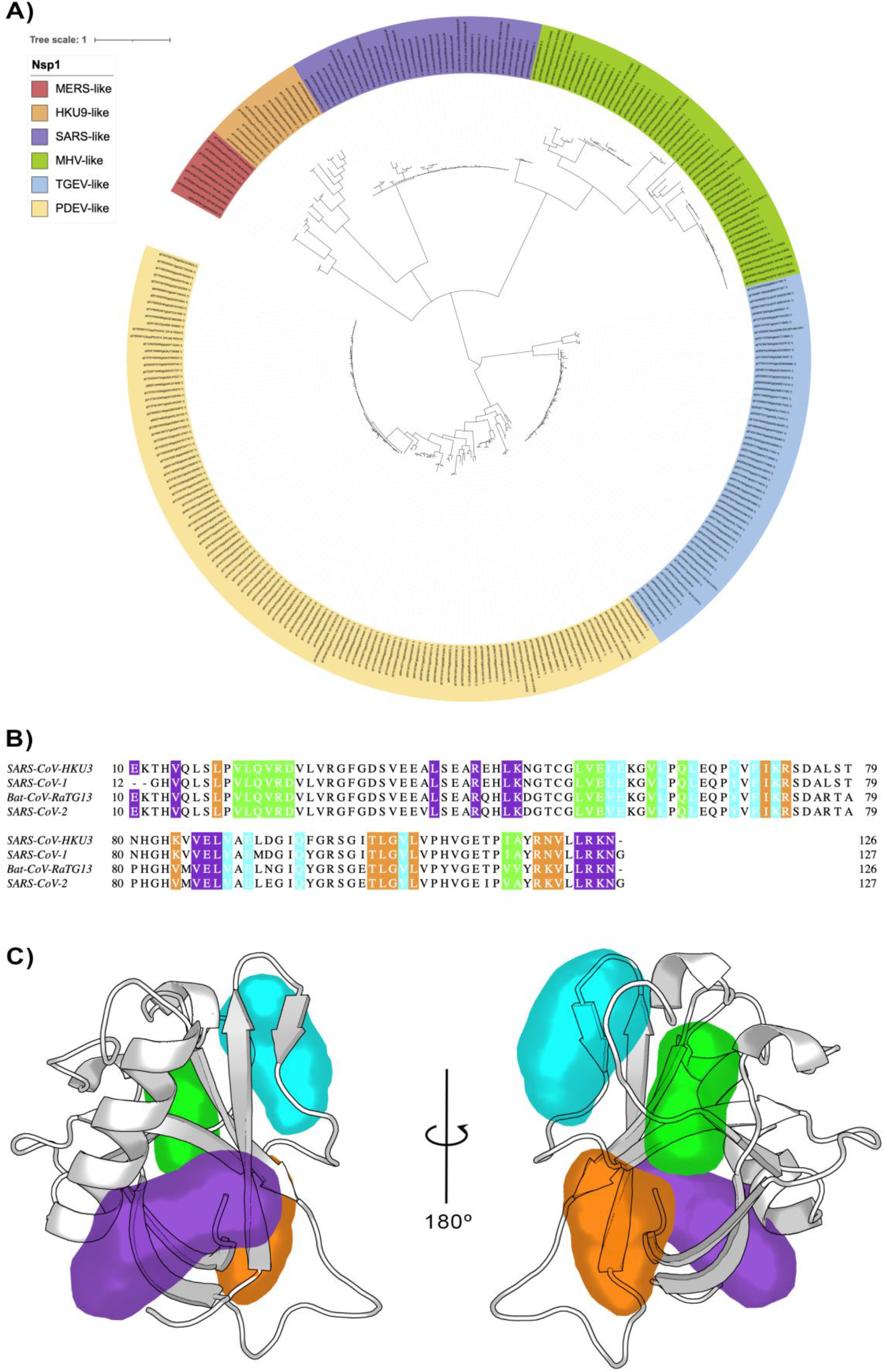
A) Phylogenetic tree based on 283 sequences from distinct *α*- and *β*-CoVs of different subgenera. The sequences were obtained from the CDD database with accession number cl41742. The scale bar indicates the number of substitutions per site in the amino acid sequence. Six different Nsp1 domain models can be identified, two for the *α*-CoVs (TGEV-like and PDEV-like), and four for the *β* CoVs genus (MERS-like, HKU9-like, SARS-like and MHV-like). B) Multi sequence alignment of the four homologues selected. The alignment was performed with the MUSCLE algorithm. ^35^ The residues of the different pockets are highlighted in the corresponding color, namely pocket 1 in purple, pocket 2 in green, pocket 3 in orange and pocket 4 in cyan. C) Representation of the pockets found in the SARS-CoV-1 Nsp1_N_ variant.

Finally, taking into account the previous data and the fact that Nsp1 proteins share a high 3D structural core identity across coronavirus species,^10,11^ we have run 1 μs of unbiased MD simulations for each of the three Nsp1 homologues considered, which show a stable conformation (Figure 9 - Figure supplement 1). Additionally, for all of them, we have evaluated the conservation of the cryptic binding pockets via extensive SWISH simulations (six replicas of 500 ns per homologue). The resulting trajectories indicate that the four pockets we reported for Nsp1_N_ of SARS-CoV-2 are conserved in the considered homologues (Figure 9C and Figure 9 - Figure supplement 2). Taken together, these results suggest that ligands binding to any of the pockets identified in SARS-CoV-2 Nsp1_N_, could also target the corresponding pocket in the evaluated homologues, ultimately paving the way to the development of a drug targeting the Nsp1_N_ of different *β*-CoVs.

## Conclusions

Nsp1 is a promising drug target for CoVs both due to its crucial role in suppressing host immune response and its sequence conservation and structural similarity across the α- and β-coronavirus families. In this paper, by using a multidisciplinary approach combining modelling, simulations, X-ray crystallography and fragment screening, we revealed druggable and partially cryptic pockets in the folded main domain of Nsp1. Our enhanced sampling simulations revealed four candidate pockets and predicted that fragments can bind to them. Not all of these are predicted to be accessible in a crystal structure, due to crystallographic contacts. The fragment screening and subsequent crystallographic structures confirm the presence of the deepest and most accessible pocket predicted by the simulations. Interestingly, the pockets are conserved across multiple coronavirus species. Moreover, we show how the fragments bound to these pockets can disrupt the Nsp1-RNA complex. Altogether, the crucial information arising from our multidisciplinary approach can provide a solid foundation for the rational drug discovery of new inhibitors not only for SARS-CoV-2, but also for other *α*- and *β*-CoVs with pandemic potential.

## Materials and Methods

### Set up of the systems

#### Preparation of Nsp1_N_

The structure of the N-terminal domain of SARS-CoV-2 Nsp1 (Nsp1_N_) was obtained from the PDB entry 7K7P, (resolution 1.77 Å), comprising residues Glu10-Asn126.^36^

#### Preparation of Nsp1_FL_

The Nsp1_FL_ structure model was obtained from AlphaFold 2.0 neural-network based structural prediction ^8,9^, which is based on the NCBI Reference Sequence: YP_009725297.1

#### Preparation of Nsp1_FL_-RNA complex

The SL1 structure corresponding to nucleotides 7-33 at the 5’ UTR of the viral mRNA was modelled with RNAComposer.^37,38^ The NCBI accession number for the reference viral RNA is NC_045512 (https://www.ncbi.nlm.nih.gov/genome/viruses/). Then, the SL1 structure obtained was docked to the Nsp1_FL_ model from AlphaFold using the HADDOCK software.^29,30^ Two docking poses of Nsp1_FL_-RNA complexes were selected based on their score and the non-covalent interactions on the interface of Nsp1_FL_ with RNA (Figure 8 - Figure supplement 1).

#### Preparation of Nsp1 in different CoV genera

The NMR structure of the SARS-CoV-1 Nsp1_N_ was obtained from PDB entry 2HSX. Since there are no experimentally determined structures for the Nsp1 of Bat CoVs HKU3 nor RatG13, we modeled their structure using Nsp1_N_ of SARS-CoV-2 (PDB entry 7K7P) as a template. The NCBI accession number for HKU3 and RatG13 is QND76032.1 and QHR63299.2, respectively, using the *automodel* function of MODELLER. ^39^

Each of these systems has been processed in the same way. First, the standard protonation state at physiological pH 7.4 was assigned to ionizable residues with the ProteinPrepare tool of the PlayMolecule server.^36^ Then, the systems were placed in a pre-equilibrated octahedral box using the four-point water model from the a99SB-disp force field,^1^ which is a modified version of TIP4P-D.^40^ The final systems considering the full-size enzyme contain the model protein, around 6,500 water molecules, and 0.15 M of NaCl, forcing the system to be neutral, leading to simulation systems comprising around 29,000-30,400 atoms for Nsp1_N_ systems, and approximately 96,400 atoms for the Nsp1_FL_-RNA complex.

All the simulations were performed using the a99SB-disp force field, which is a modified form of the a99SB force field that improves the modeling of intrinsically disordered peptides while retaining the accurate description of folded proteins. To parameterize the SL1 from RNA, the RNA-Shaw force field was used.^41^

### Molecular Dynamics simulations

#### Unbiased Molecular Dynamics simulations of Nsp1_N_

All the atomistic MD simulations were performed using the GROMACS 2021.3 package^42^ employing the a99SB-disp force field.^43^ Energy minimization was conducted using 50,000 steps of the steepest descent algorithm and setting the tolerance to 100 kJ mol^-1^nm^-1^. The equilibration process was performed in two steps, applying harmonic restraints to all heavy atoms in the system (harmonic constant: 1000 kJ mol^-1^nm^-1^). Firstly, a 5 ns heating in the NVT ensemble was performed, using the V-rescale (τ = 0.1 ps) as thermostat.^44^ Two different groups were used for temperature coupling: one for the protein and a second one comprising water molecules and ions. The reference temperature was set to 310 K. Secondly, the system was equilibrated during 15 ns in the NPT ensemble using V-rescale^44^ (τ = 0.5 ps) and Berendsen^45^ (τ = 0.5 ps) as thermostat and barostat, respectively. The same temperature coupling groups were kept during the NPT equilibration step. The final structure from the equilibration process was used as a starting point for the MD simulations. All systems were simulated with periodic boundary conditions using an NPT ensemble using the same parameters as in the equilibration step and removing the harmonic restraints. The Particle Mesh Ewald (PME) method was used for treating long-range electrostatics using a cutoff of 12 A.^46^ A time step of 2 fs was used for all simulations after imposing constraints on the hydrogen stretching modes. We ran one replica of 1 μs of Nsp1_N_ from SARS-CoV-2, and one replica of 1 μs of the three Nsp1_N_ homologues (human SARS-CoV-1, and two Bat CoVs). For the Nsp1_FL_-RNA complex we ran 1 replica of 500 ns for each model, A and B. Considering all the systems simulated leads to a total simulation time of 5 μs.

### SWISH simulations

SWISH (Sampling Water Interfaces through Scaled Hamiltonians)^4,11^ is a Hamiltonian replica exchange enhanced sampling technique that increases the conformational sampling of proteins by scaling the interaction of the apolar atoms of the protein with the water molecules. In this way, water molecules acquire more hydrophobic physicochemical properties that allow them to induce the opening of hydrophobic cavities. Including organic fragments in the solvent during the SWISH simulations has been shown to stabilize the cavities that open up during the simulation.^24,47^ All SWISH simulations presented in this work, i.e., of the Nsp1_N_ of SARS-CoV-2 and the three CoV homologues, were run employing the same protocol: six different parallel 500-ns replicas, each one at a specific scaling factor (λ, ranging evenly from 1.00 to 1.35) value, in the presence of benzene (1 M concentration) as co-solvent. Since we ran six replicas per SWISH simulation, the total accumulated time is 12 μs. Besides the scaling factors, all other parameters for both equilibration and production are the same as in the ones used for the unbiased MD simulations. Before each production run, six independent equilibration steps were carried out, one for each λ value. A contact-map-based bias was introduced for each replica to prevent the possible unfolding of the protein in high scaling factor replicas. The optimal upper wall value for the contact map was tuned based on the unbiased simulations. The benzene molecules for the mixed-solvent simulations were parametrized using Gaussian16^48^ with Amber GAFF-2 force field^49^ and RESP charges.

### Pocket detection

#### Crystal structures

In order to evaluate whether binding pockets exist in the Nsp1 and quantify their physicochemical properties, the crystal structure of Nsp1_N_ (PDB entry 7K7P) was analyzed with Fpocket^12^ and DoGSiteScorer that is available as part of the modelling server ProteinPlus.^21^ Fpocket is a geometry-based cavity detection algorithm that employs Voronoi tessellation and α spheres to identify pockets in the protein structure. In this context, an α sphere is defined as a sphere that contacts four atoms on its boundary and contains no internal atom. Similarly, DoGSiteScorer is an algorithm for pocket and druggability prediction that employs 3D difference of gaussian filters to detect cavities and a support vector machine to score the identified binding sites.

#### MD simulations

The trajectories were analyzed with MDpocket,^31^ an open source to detect binding pockets along MD simulations. MDpocket was run over down-sampled and reference-aligned trajectories with a time step of 100 ps between each frame. The corresponding minimized Nsp1_N_ structure was chosen as a reference structure for each different system. The outputs let us identify and visualize the pockets observed throughout the whole simulation time with the PyMol software.^50^ This analysis has been performed along all the trajectories resulting from MD and SWISH simulations. Additionally, the software lets us compute the volume of the selected pockets.

### Druggability assessment

*FTDyn and FTMap algorithms* proved to be accurate in locating binding hot spots in proteins, i.e., regions of the surface that majorly contribute to the free energy of binding. This approach is based on the fast and accurate distributions of 16 different small organic probes on the protein surface. Each probe is docked billions of times and map onto the target surface and scored according to an energy-based function.^51^ Hence the lowest energy probes are retained, locally minimized, and clustered. Ultimately, clusters of different probes are clustered together into a consensus cluster. The main binding hot spot is then identified as the consensus cluster containing the highest number of different probe fragments. We first employed FTDyn, a faster version of the FTMap algorithm without local minimization, to identify the most bound-like conformation in our unbiased MD simulations. We extracted 25 representative conformations from the MD simulation of SARS-CoV-2 Nsp1_N_, one from each of the first 25 most populated clusters. The 10000-frames trajectory was clustered employing the *gromos* algorithm with a cut-off of 0.1 nm. Hence, we processed these structures with FTDyn server and determined the median number of interactions between probe molecules and each protein residue across the structural ensemble. We then assigned a contact score to each structure in the ensemble. The contact score is calculated as the sum of the number of residues in a given structure with a number of contacts higher than the median. We then calculated an ensemble score averaging over the 25 contact scores of the structures in the ensemble. Finally, we retained the structures in our 25-structures ensemble with a contact score higher than the average, as they were considered to be the best approximations of bound-like conformations. We further processed these 11 structures with FTMap to have a better prediction of binding hot spots on the Nsp1_N_ surface.

### Crystallographic data

The N-terminal domain of SARS-CoV2 Nsp1 was purified and crystallised as previously described (add ref). 60 potential fragment hits obtained from computational fragment screening of the Maybridge Ro3 library were purchased and fragments soaked into Nsp1 crystals and validated through X-ray diffraction experiments in quasi-automated mode at ESRF beamline MASSIF-1. Data analysis of the 60 data sets was conducted in the multi-crystal system PanDDA.^52^ One hit was subsequently verified by manual inspection in COOT followed by refinement in Phenix.

### Phylogenetic analysis

The 283 sequences of different *α*- and *β*-CoVs of different subgenera were obtained from the CDD database, family accession number cl41742. The sequences were aligned with MEGA, version 11.0.11.^53^ The resulting multi sequence alignment was used to construct the Maximum-Likelihood Tree with MEGA. The resulting tree was rendered with the iTol web server.^54^

## Acknowledgements

We acknowledge PRACE and the Swiss National Supercomputing Centre (CSCS) for large supercomputer time allocations on Piz Daint, project IDs: pr126, s1107. FLG acknowledges the Swiss National Science Foundation and Bridge for financial support [projects number: 200021_204795 and 40B2-0_203628]. We thank Dr. Matthew Bowler at ESRF beamline Massif-1 for his excellent support.

## Author contributions

Alberto Borsatto, conceptualization, methodology, validation, visualization, writing – original draft. Obaeda Akkad, Methodology, visualization.

Ioannis Galdadas, conceptualization, methodology, validation, visualization, supervision. Shumeng Ma, methodology, validation.

Shymaa Damfo, methodology, validation.

Shozeb Haider, conceptualization, supervision.

Frank Kozielski, conceptualization, supervision.

Carolina Estarellas, conceptualization, validation, visualization, supervision, writing – original draft.

Francesco Luigi Gervasio, conceptualization, supervision, funding acquisition.

All the authors have contributed to the writing - review and editing.

## Supportive information

**Figure 2 - Figure supplement 1.**
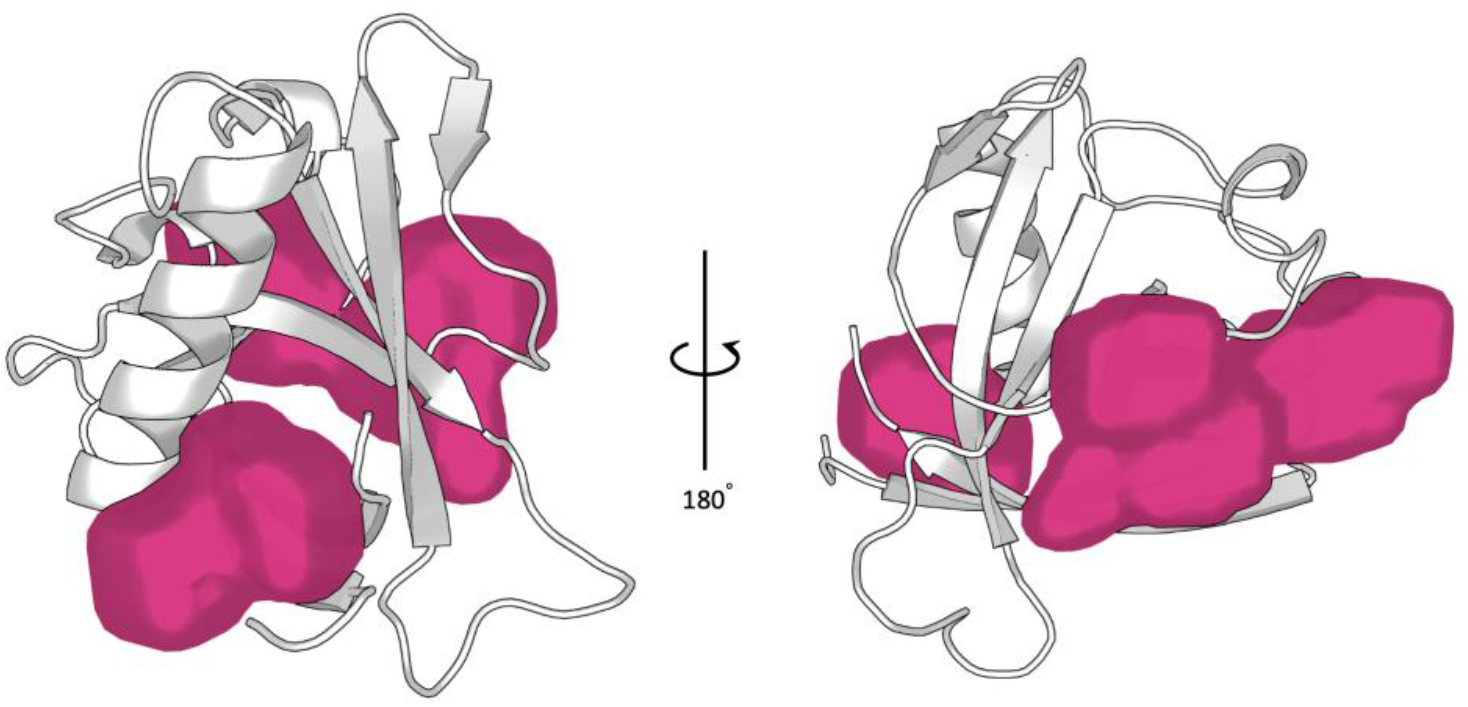
Cavities identified by the Fpocket software on the Nsp1_N_ crystal structure (PDB entry 7K7P).^22^

**Figure 3 - Figure supplement 1.**
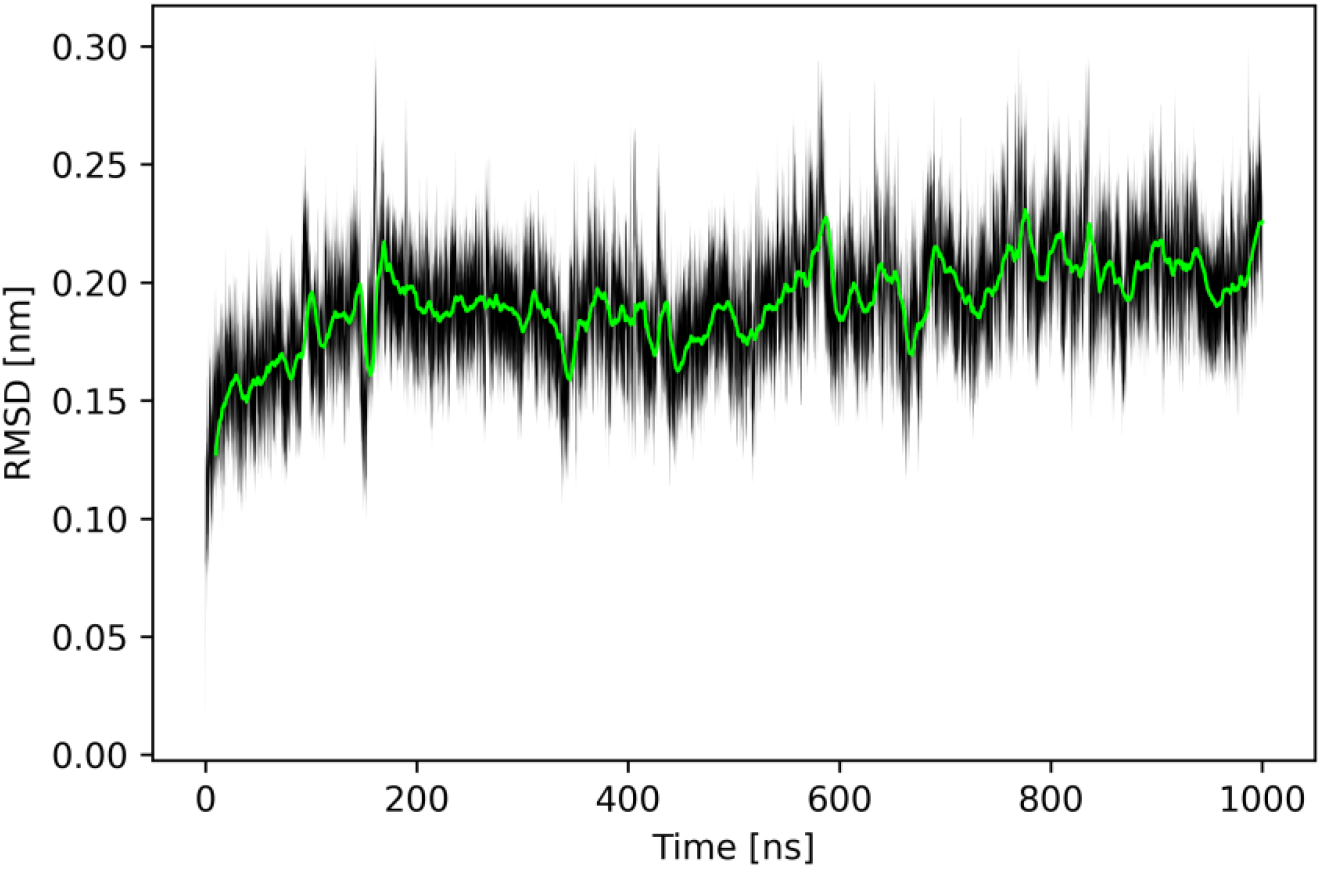
Time average of the RMSD for Nsp1_N_. The black shaded area delimits the RMSD values within two standard deviations from the time average.

**Figure 4 - Figure supplement 1.**
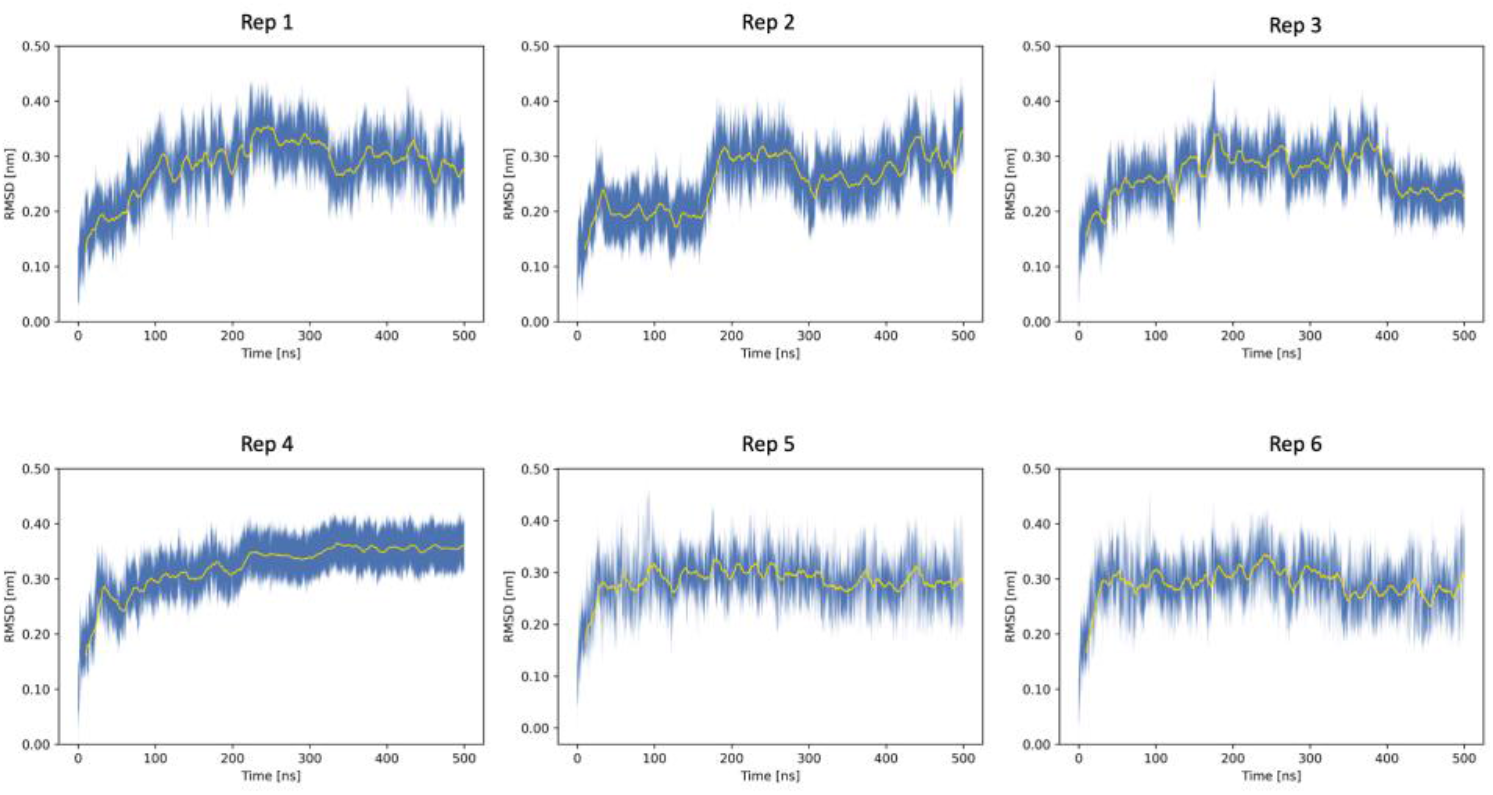
Time averages of the RMSD for Nsp1_N_ across the six swish replicas. The blue shaded area delimits the RMSD values within two standard deviations from the time average.

**Figure 4 - Figure supplement 2.**
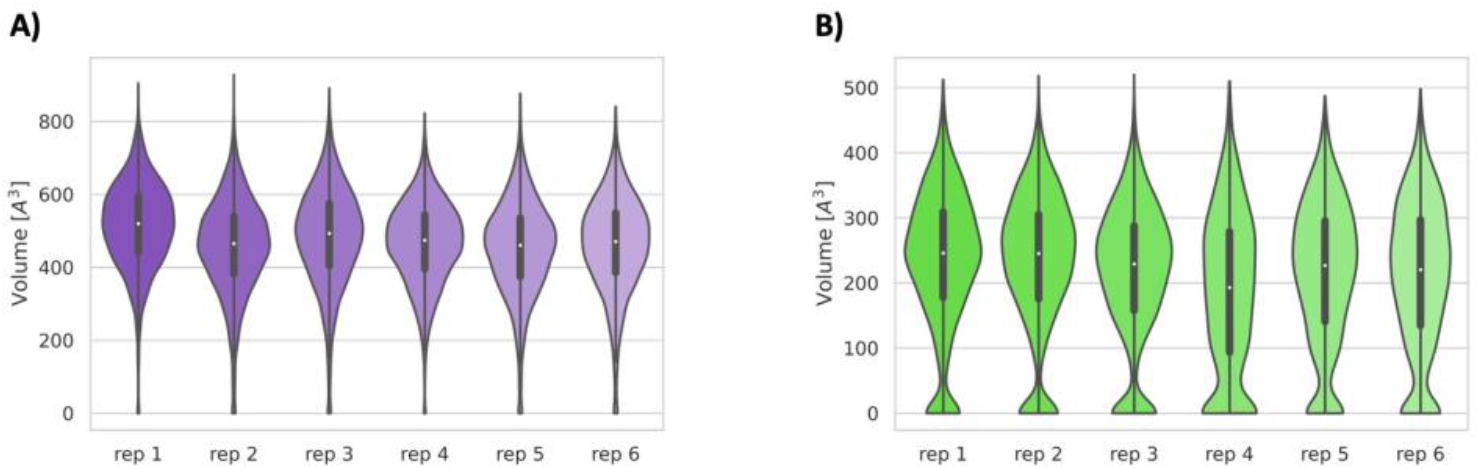
Volume distributions of the cryptic binding sites pocket 1 (A) and pocket 2 (B) along the six replicas of the SWISH simulation of SARS-CoV-2 Nsp1_N_.

**Figure 8 - Figure supplement 1.**
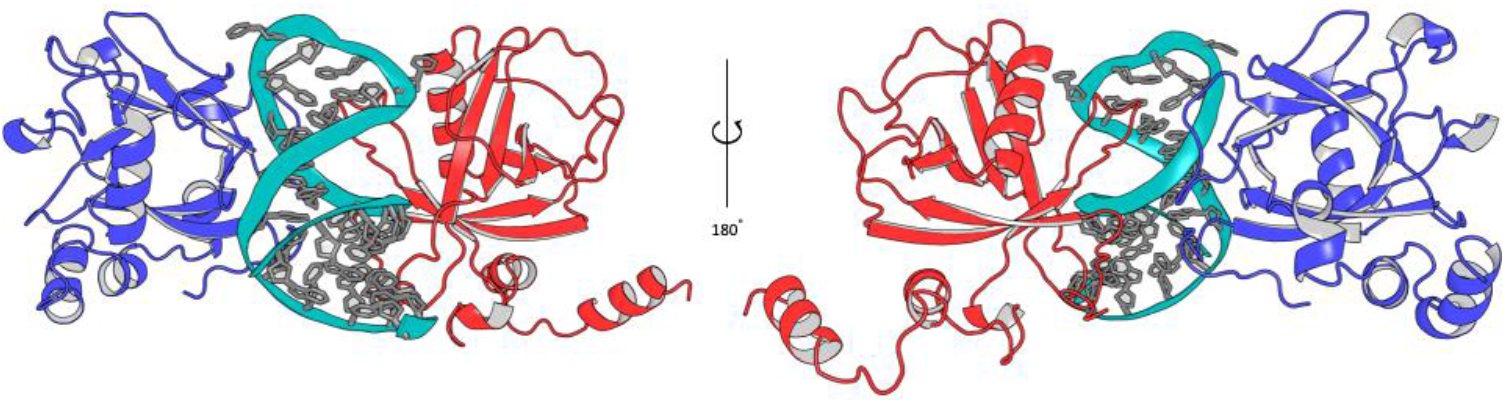
Different Nsp1_FL_-RNA models proposed in this work. The Nsp1_FL_ from model A (blue) and B (red) are represented.

**Figure 8 - Figure supplement 2.**
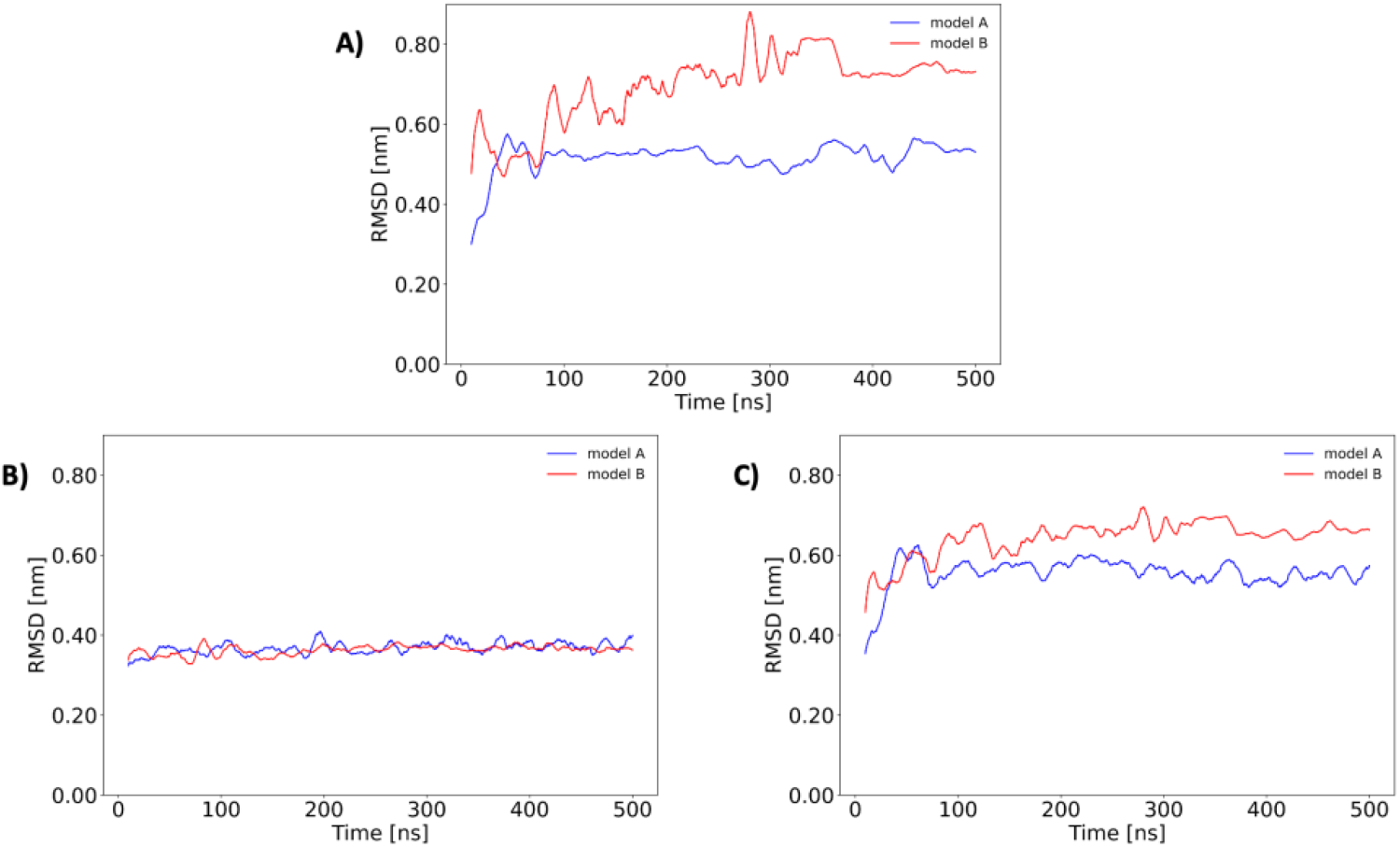
Root mean square deviation (RMSD, nm) along the 500 ns of unbiased MD simulations considering A) the backbone atoms of the Nsp1_FL_ protein from the Nsp1_FL_-RNA complex, B) only the RNA atoms of the SL1 from the complex and (C) the backbone atoms of Nsp1_FL_ and all the RNA atoms of the full Nsp1_FL_-RNA complex.

**Figure 9 - Figure supplement 1.**
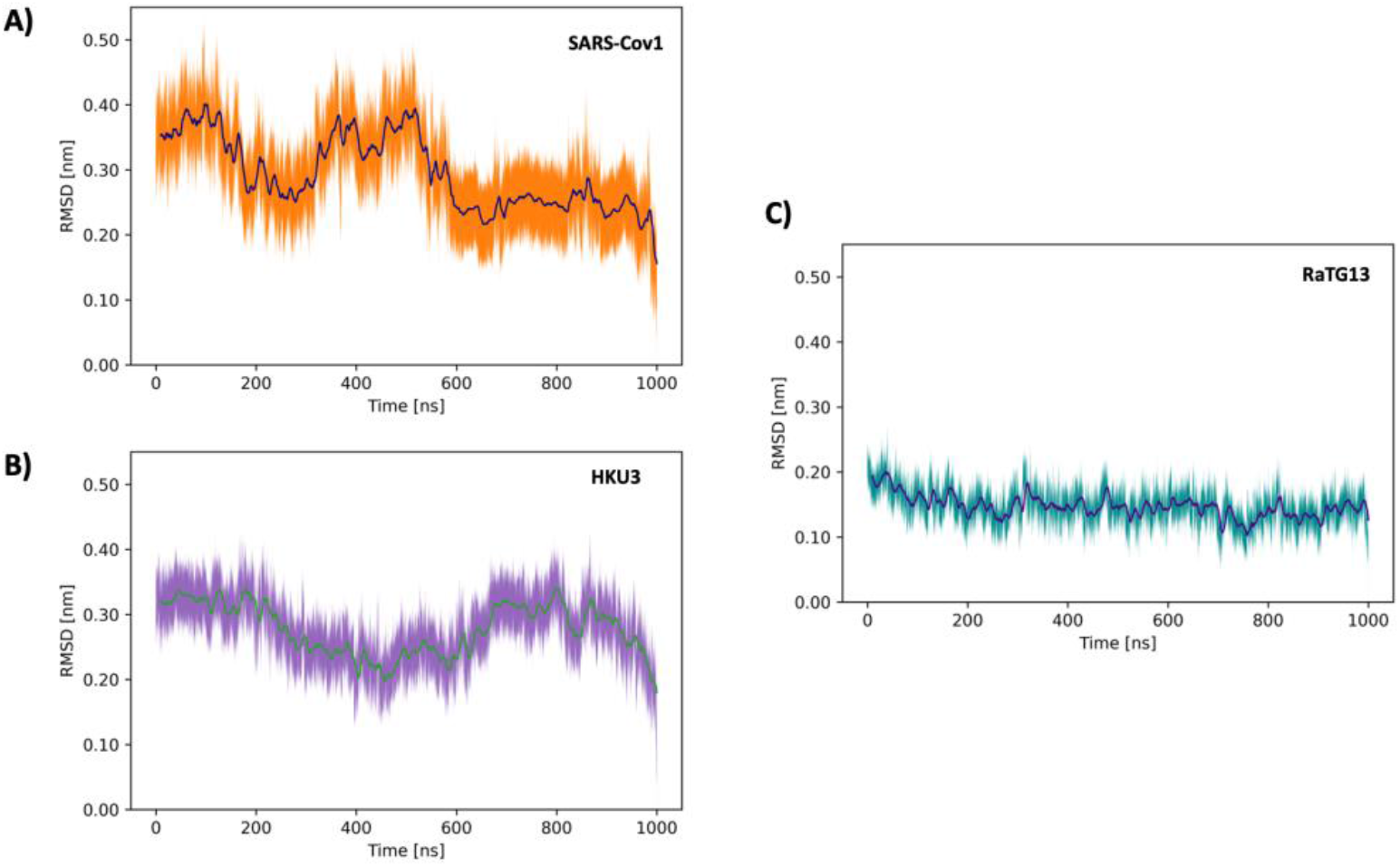
Time averages of the backbone RMSD along the 1 μs unbiased MD simulation for Nsp1_N_ of A) SARS-CoV1, B) SARS-HKU3 and (C) Bat-CoV-RatG13. The shaded area delimits the RMSD values within two standard deviations from the time average.

**Figure 9 - Figure supplement 2.**
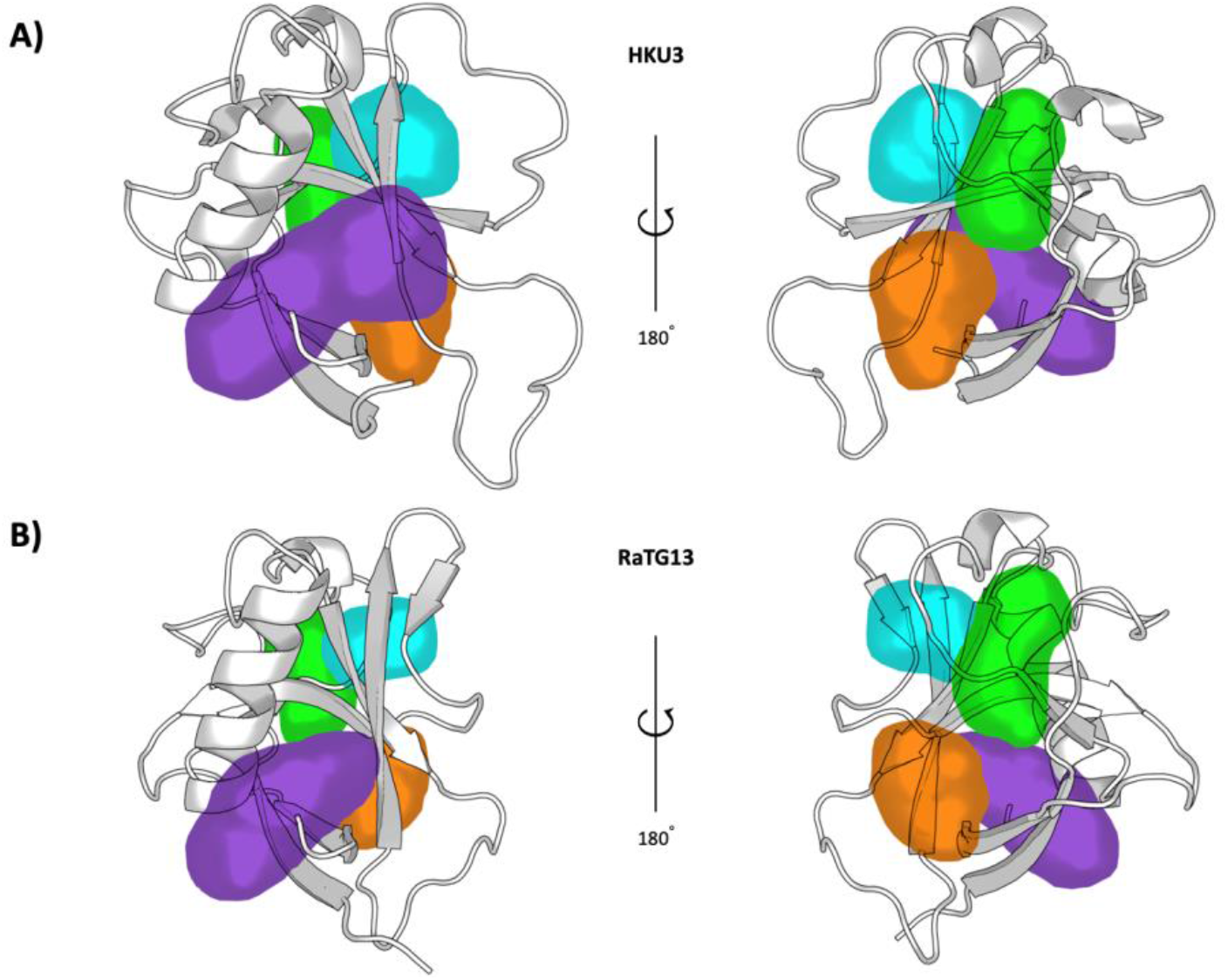
Representation of the pockets found in the Bat-CoV-HKU3 (A) and Bat-Cov-RaTG13 (B) Nsp1_N_ variants.

**Figure 5 – Table supplement 1.** Consensus clusters obtained from the FTMap program.

| Selected structures | # Consensus clusters | # Of fragments in the top 3 consensus clusters | Pocket location of the top 3 consensus clusters |
| --- | --- | --- | --- |
| 1 | 7 | 24 | Pocket 1 |
|  |  | 17 | Pocket 1 |
|  |  | 16 | Pocket 3-4 |
| 2 | 8 | 22 | Pocket 3-4 |
|  |  | 17 | Pocket 1 |
|  |  | 16 | Pocket 1 |
| 3 | 12 | 17 | Pocket 1 |
|  |  | 17 | Pocket 3 |
|  |  | 14 | Pocket 1 |
| 4 | 7 | 22 | Pocket 1 |
|  |  | 18 | Pocket 3-4 |
|  |  | 16 | Pocket 1 |
| 5 | 11 | 20 | Pocket 1 |
|  |  | 14 | Pocket 3-4 |
|  |  | 13 | Pocket 1 |
| 6 | 11 | 16 | Pocket 1 |
|  |  | 14 | Pocket 3 |
|  |  | 11 | Pocket 2 |
| 7 | 10 | 16 | Pocket 1 |
|  |  | 16 | Pocket 1 |
|  |  | 15 | Pocket 4 |
| 8 | 11 | 16 | Pocket 1 |
|  |  | 15 | Pocket 3-4 |
|  |  | 13 | Pocket 1 |
| 9 | 8 | 15 | Pocket 1 |
|  |  | 19 | Pocket 1 |
|  |  | 16 | Pocket 3-4 |
| 10 | 11 | 18 | Pocket 1 |
|  |  | 16 | Pocket 1 |
|  |  | 15 | Pocket 1 |
| 11 | 11 | 25 | Pocket 1 |
|  |  | 16 | Pocket 1 |
|  |  | 15 | Pocket 2 |

**Figure 6 – Table supplement 1.** Data collection, data processing, and model refinement statistics for the SARS CoV-2 Nsp1-2E10 complex. Data in parenthesis correspond to the highest resolution shell.

|  | **Nsp1-2E10** |
| --- | --- |
| **PDB ID** | xxx |
| **Wavelength [Å]** | 0.96546 |
| **Resolution range [Å]** | 32.51 - 1.10 (1.12 - 1.10) |
| **Space group** | P4<sub>3</sub>2<sub>1</sub>2 |
| **Unit cell parameters (Å, °)** | $a=b=36.688$, $c=140.928$,<br>$\alpha=\beta=\gamma=90$ |
| **Total reflections** | 464682 (12250) |
| **Unique reflections** | 40443(1953) |
| **Multiplicity** | 11.5 (6.3) |
| **Completeness [%]** | 99.9 (98.1) |
| **Mean I/sigma(I)** | 18.4 (2.3) |
| **R-meas** | 0.067 (0.750) |
| **R-pim** | 0.020 (0.291) |
| **CC<sub>1/2</sub>** | 0.998 (0.801) |
| **R<sub>cryst</sub>/R<sub>free</sub> [%]** | 19.8 (24.2) / 22.3 (23.8) |
| **Total no. of non-hydrogen atoms (protein)** | 1082 |
| **No. of protein/ligand/solvent atoms** | 952 / 26 / 104 |
| **Average B-factor/protein/ligands/solvent** | 19.6 / 18.6 / 31.1 / 26.2 |
| **RMSD (bonds, angles)** | 0.012 / 1.33 |
| **Ramachandran**<br>**favored/allowed/outliers/rotamer outliers [%]** | 98.2 / 1.8 / 0/0 / 0/0 |
| **Clashscore** | 3.6 |

**Figure 9 – Table supplement 1.** Pairwise identity percentages of selected Nsp1 sequences for representative *α*- and *β*-CoVs sub-families. NCBI accession numbers the sequences used for the analysis are as follows: TGEV *6IVC_A*, PDEV *5XBC_A*, SARS CoV 1 *NP_828860.2*, SARS CoV 2 *YP_009725297.1*, HKU9 *P0C6T6*, MERS *YP_009047229* and MHV *YP_209244*.

| **CoV-like** | *SARS-1* | *SARS-2* | *HKU9* | *MERS* | *MHV* | *TGEV* | *PDEV* |
| --- | --- | --- | --- | --- | --- | --- | --- |
| *SARS-1* | 100 |  |  |  |  |  |  |
| *SARS-2* | 86,1 | 100 |  |  |  |  |  |
| *HKU9-* | 20,9 | 19,5 | 100 |  |  |  |  |
| *MERS-* | 20,9 | 17,1 | 24,5 | 100 |  |  |  |
| *MHV-* | 16,9 | 16,4 | 4,1 | 18 | 100 |  |  |
| *TGEV-* | 17,4 | 16,9 | 13,3 | 7,9 | 9 | 100 |  |
| *PDEV-* | 17,9 | 16,1 | 15,2 | 8,7 | 11,5 | 18,2 | 100 |

